# The *Drosophila* Circadian Clock Gene *Cycle* Controls the Development of Clock Neurons

**DOI:** 10.1101/2023.10.29.564626

**Authors:** Grace Biondi, Gina McCormick, Maria P. Fernandez

**Affiliations:** Department of Neuroscience and Behavior, Barnard College. New York, NY 10027

**Keywords:** neuronal development, fasciculation, circadian clock, cycle, *Drosophila*

## Abstract

Daily behavioral and physiological rhythms are controlled by the brain’s circadian timekeeping system, a synchronized network of neurons that maintains endogenous molecular oscillations. These oscillations are based on transcriptional feedback loops of clock genes, which in *Drosophila* include the transcriptional activators *Clock (Clk)* and *cycle (cyc)*. While the mechanisms underlying this molecular clock are very well characterized, the roles that the core clock genes play in neuronal physiology and development are much less understood. The *Drosophila* timekeeping center is composed of ∼150 clock neurons, among which the four small ventral lateral neurons (sLN_v_s) are the most dominant pacemakers under constant conditions. Here, we show that downregulating the clock gene *cyc* specifically in the *Pdf*-expressing neurons prevents leads to decreased fasciculation both in larval and adult brains. This effect is due to a developmental role of *cyc*, as both knocking down *cyc* or expressing a dominant negative form of *cyc* exclusively during development lead to defasciculation phenotypes in adult clock neurons. *Clk* downregulation also leads to developmental effects on sLNv morphology, although *cyc* and *Clk* manipulations produce distinct phenotypes. Our results reveal a non-circadian role for *cyc*, shedding light on the additional functions of circadian clock genes in the development of the nervous system.

## Introduction

The proper wiring of neuronal circuits during development is essential for the neuronal control of behavior. Across animal species, sleep/wake cycle rhythms, as well as many other behavioral and physiological rhythms, are controlled by the circadian timekeeping system, a network of neurons that maintains endogenous molecular oscillations and rhythmic behavior with a ∼24 hour period ^1^. The proper functioning of this circadian network requires the formation of synaptic and peptidergic connections during development ^2,3^.

The *Drosophila* circadian clock neuron network comprises ∼150 neurons and is the functional equivalent of the mammalian suprachiasmatic nuclei, which contain 20,000 neurons in mice^4–6^. All circadian clock neurons contain an intracellular molecular clock consisting of a transcriptional feedback loop of clock genes^7^. CLOCK (CLK) and CYCLE (CYC) are heterodimeric transcriptional activators that directly activate transcription of the *period* (*per*) and *timeless* (*tim*) genes. PER and TIM encode repressors that inhibit CLK-CYC function. Subsequently, PER and TIM are degraded, which enables the cycle to reinitiate every morning. CLK and CYC also interact with other genes in a secondary circadian loop by activating the genes *vrille* (*vri*), and *Par domain proteinε* (*Pdp1ε*)^8,9^. *Clk* and *cyc* expression can be detected in almost all clock neurons even before some of these neurons show molecular oscillations^10^, suggesting that these genes serve functions that precede the establishment of molecular rhythms.

*Drosophila* clock neurons are classified into multiple clusters with distinct patterns of gene expression, anatomy, physiology, and synaptic connectivity^5,6,11–16^. Among these clusters, the small ventral lateral neurons (sLN_v_s) are considered the most dominant pacemakers since they are critical for maintaining behavioral rhythmicity under constant darkness and temperature (DD, or free-running)^17–20^. The sLN_v_s release the neuropeptide Pigment Dispersing Factor (PDF)^21^, a key output signal within the clock neuron network ^22^. PDF accumulates rhythmically at the sLN_v_ dorsal termini^23^ and can be released from both the neurites and soma^24^. Loss of PDF severely reduces the amplitude of the endogenous circadian rhythm and shortens its free-running period in DD^21^. The large LN_v_s also produce PDF but do not play a role in maintaining rhythms in DD^17^.

The projections of the four sLN_v_s form a bundle and remain fasciculated as they extend from the ventral to the dorsal brain during development. These four projections are usually difficult to distinguish from each other until they begin to defasciculate in the dorsal protocerebrum^25^ and extend their dorsal arborizations toward the area where dorsal clusters of clock neurons are located^26^. In adult flies, the dorsal termini of the sLN_v_ projections show rhythmic structural plasticity^27^, which relies on daily and circadian rhythms in outgrowth and fasciculation^28–31^. Both *Clk* and *cyc* mutants have lower *Pdf* RNA levels, and the PDF peptide can barely be detected in the sLNv projections^23,32^.

*Cyc* is a homolog of the mammalian gene *Bmal1*, although CYC protein levels do not cycle, unlike BMAL1 and several other *Drosophila* circadian proteins^33^. There is growing evidence for non-circadian functions of BMAL1. First, its downregulation induces apoptosis and cell-cycle arrest in Glioblastoma Stem Cells (GSC), and it was found to preferentially bind metabolic genes and associate with active chromatin regions in GSCs^34^. Second, brain knockdown of *Bmal1* using CRISPR/Cas9 made glioblastomas grow at faster rates than controls^35^, and similar effects were observed in B16 melanoma cells. Moreover, *Bmal1*(-/-) mice exhibit defects in short- and long-term memory formation^36^ and show reduced lifespan and multiple symptoms of premature aging ^37^. Overall, results from studies in different animal models suggest that *Bmal1* plays a role in the development of various neurological disorders^38^.

The *Drosophila* sLN_v_s offer an excellent model for exploring the non-circadian roles of canonical clock genes such as *cycle*. To determine if the phenotypes previously observed for *cyc* mutants are specific to PDF expression or involved a broader, non-circadian effect in the development of PDF-expressing cells, we downregulated *cyc* specifically in the *Pdf*-expressing cells and observed pronounced defasciculation of the sLN_v_ projections. Similar phenotypes were observed upon expression of a dominant negative form of *cyc*. Moreover, we found that *cyc* downregulation in *Pdf*+ cells during development is sufficient to prevent the fasciculation of the adult sLN_v_s and results in the loss of behavioral rhythms in adult flies. Manipulations of *Clk* expression also affect sLNv morphology, although remarkably, the phenotypes of *Clk* and *cyc* manipulation differ. Our results show that *cyc* plays a role in the development of pacemaker neurons, which is likely independent of its role in the circadian molecular oscillator.

## Results

### *cyc* downregulation in circadian pacemaker neurons affects the formation of sLN_v_ axon bundles

Mutations in both *Clk* and *cyc* severely reduce *pdf* RNA and neuropeptide levels^23,32^. In *cyc* null mutants, sLN_v_s projections cannot be detected in larval or adult brains stained with PDF antibodies^23,39,40^. We observed that *cyc* null mutants (*cyc^01^*) showed a substantial reduction in PDF levels at ZT2 compared to Canton-S controls, consistent with previous studies (Fig. 1B). However, we also noticed the presence of thin, misrouted sLN_v_ projections in *cyc^01^* flies at higher magnification and intensity (Fig. 1B, right panels). Upon close observation, PDF could often be detected in the sLN_v_ projections. However, these projections did not form the stereotypical bundle observed in control brains when extending from the anterior medulla toward the dorsal area of the brain.

**Figure 1.**
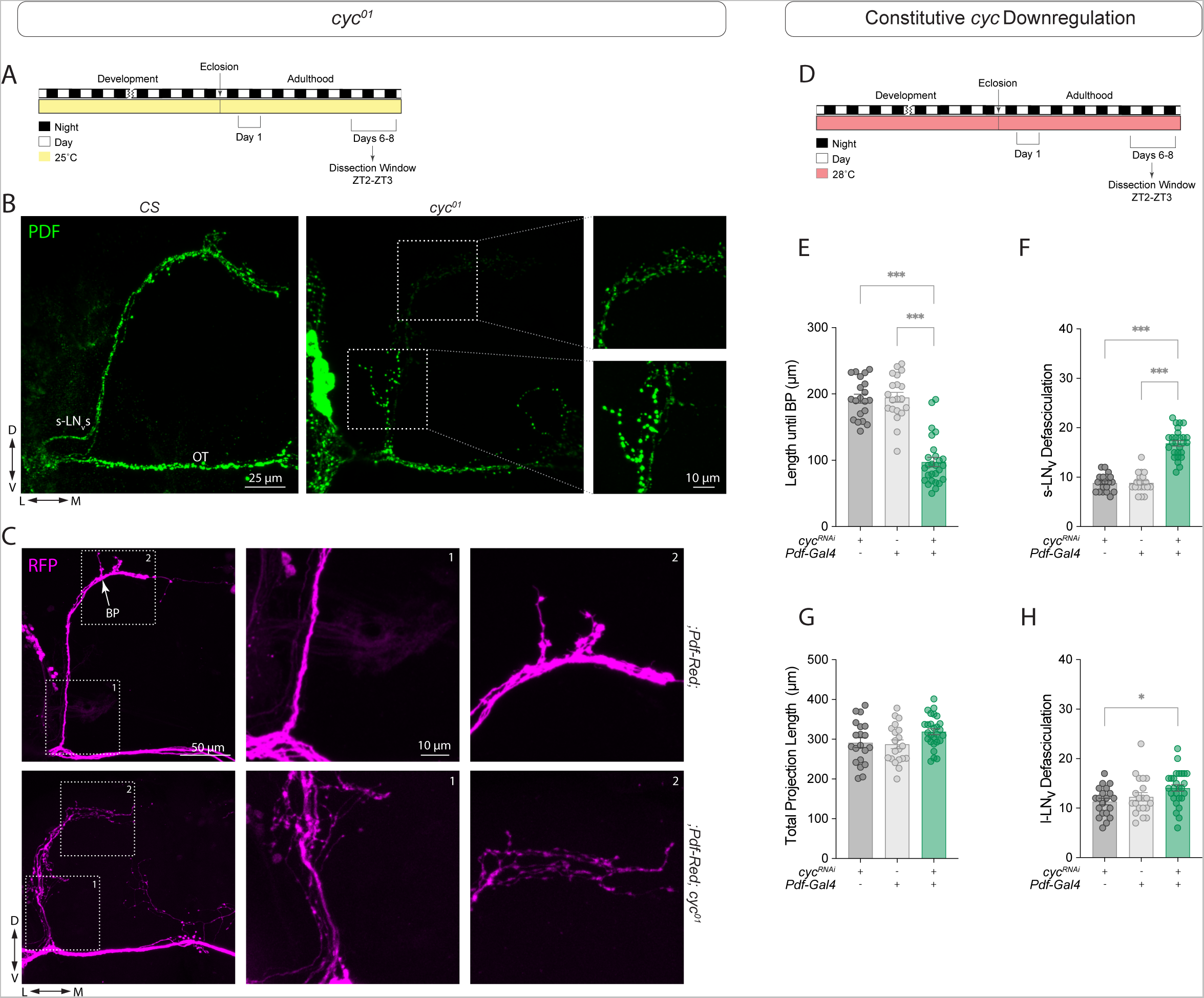
*Cyc* downregulation in circadian pacemaker neurons prevents the formation of sLN_v_s axon bundles. The *cyc^01^* mutant has disrupted sLN_v_ morphology. (A) Representative timeline of the *cyc^01^* mutant experiments in the figure. Flies were kept at 25°C for their entire lifespan. Experiments were then performed within Days 6-8 post-eclosion from pupae. (B) Representative confocal images of anti-PDF staining in Canton-S control and *cyc^01^*adult male brains. The sLN_v_s and optic tract (OT) are indicated. Scale bar = 25 µm. The inserts on the right show the sLN_v_ projections with the signal intensity adjusted for visibility in the *cyc^01^*mutants. The top insert shows the distal (dorsal) area and the bottom insert shows the proximal (ventral) area of the sLN_v_ projections. Scale bar = 10 µm. (C) Representative confocal images of *Pdf-RFP* controls and *;Pdf-RFP;cyc^01^* experimental flies stained with anti-RFP (magenta). In the control brains, the stereotypical branching point (BP) of the dorsal projections is indicated. Scale bar = 25 µm. Boxes with dashed lines indicate the proximal (1) and distal (2) projections, corresponding to the labeled projection images in the center and right panels, respectively. Scale bar = 10 µm. Downregulation of *cyc* in the *Pdf+* neurons result in morphology phenotypes. (D) Representative timeline of the *cyc^01^* mutant experiments in the figure. Flies were kept at 28°C for their entire lifespan. Experiments were then performed within Days 6-8 post-eclosion from pupae. (E-H) Quantification of the LN_v_ morphology phenotypes of experimental flies in which a *cyc^RNAi^* transgene was driven by a ;*Pdf-RFP,Pdf*-*Gal4*;*Tub-Gal80^ts^* driver compared to the parental controls. The sLN_v_ projection length until the branching point (BP) (E), the total number of intersections of the sLN_v_ ventral projections (F), the total sLN_v_ projection length (G), and the total number of intersections of the lLN_v_ projections along the optic tract (OT) (H) are shown. Graphs are representative of three independent experiments, with each dot representing one brain. For each genotype, across all experiments, the number of subjects (n) fall in the range: 20 ≤ n ≤ 27 Statistical comparisons were done with one-way ANOVAs followed by Tukey post hoc tests. Differences that are not significant are not indicated. *p < 0.05, *** p < 0.001. Error bars indicate SEM.

Because highly defasciculated projections might contribute to the weaker PDF levels observed in *cyc^01^*mutants, we examined the structure of the sLN_v_ projections using a *Pdf*-RFP transgene, in which a cytosolic Red Fluorescent Protein (RFP) is controlled by the *Pdf* regulatory sequence^41^. In control brains, the projections from the four sLN_v_s remain fasciculated, forming a bundle until reaching the superior medial protocerebrum (SMP). In contrast, the sLN_v_s of *cyc^01^* mutants often began to defasciculate in the ventral brain, near their cell bodies (Fig. 1C). Some sLN_v_ projections were severely misrouted and did not reach the dorsal brain, extending instead toward the midline or other brain regions. As a result, it was possible to distinguish individual projections from each sLN_v_ even in the ventral brain in *cyc^01^* mutant brains. This is almost never observed in control brains until the sLNv projections reach the main branching point in the SMP.

To test whether the effect of *cyc* loss on the LN_v_s is cell-autonomous, we next expressed a UAS-*cyc* dsRNA transgene (UAS-*cyc^RNAi^*) using the *Pdf-Gal4* driver. We quantified the length of the projections, starting at the point where the projections of the sLN_v_s intersect with those of the lLN_v_s (“point of origin”, POI, Fig. S1A), until the first branching point (“branching point”, BP, Fig. S1A). This branching point is where the sLN_v_s ramify and extend their stereotypical arborizations in the dorsal protocerebrum in control brains, and these arborizations show daily, clock-controlled rhythms in their fasciculation and outgrowth^27^. Downregulating *cyc* in the *Pdf*-expressing cells significantly decreased the distance to the branching point (Fig. 1E). This phenotype resembled a milder version of the *cyc^01^* phenotype, since defasciculation occurred shortly after the projections begin to extend from the cell bodies. Using a modified Scholl’s analysis^42^, we quantified the degree of branching in the projections starting at the POI. We observed pronounced defasciculation in the sLN_v_ projections in *Pdf* > *cyc^RNAi^*flies (Fig. 1F). The total projection length was not different from that of the controls (Fig. 1G), and the defasciculation phenotype was not observed in the contralateral projections that extended from the lLN_v_s (Fig. 1H). Since the lLN_v_s are born later in development during metamorphosis^25^, this result suggests that *cyc* plays a role in neuronal development during an earlier developmental stage, when the sLN_v_s begin to extend their projections toward the dorsal brain.

Expression of dominant negative forms of *Clk* and *cyc* is an effective strategy for preventing circadian molecular oscillations in specific groups of clock neurons^43–45^. In these dominant negative forms, the DNA binding ability is disrupted while the ability to heterodimerize is preserved^43^. Based on the phenotypes induced by *cyc* downregulation, we asked if expressing a dominant negative form of *cyc* in the sLN_v_s also leads to aberrant projection morphology. We found that sLN_v_s expressing Δ-*cyc* using *Pdf*-*Gal4* had a significantly shorter distance until the branching point (Fig. S2B-C) and a greater degree of sLN_v_ projection defasciculation (Fig. S2B,D), similar to the effects observed in *Pdf* > *cyc^RNAi^* flies. The total projection length and the projections of the lLN_v_s were unaffected (Fig. S2E,F).

### PER levels in *Pdf*+ neurons are reduced upon cell-specific *cyc* knockdown

CYC activates *per* transcription, and thus, PER levels in the brain are significantly reduced in *cyc* mutants^33^. To test whether the phenotypes of *cyc^RNAi^* expression in the *Pdf*-expressing neurons are consistent with what would be expected from *cyc* downregulation, we compared PER levels in the parental control (*Pdf-Gal4, Pdf-RFP/+*) with those in *Pdf > cyc^RNAi^*flies at the end of the night (ZT23), when PER nuclear levels are highest^46^. We found that nuclear PER levels in *Pdf > cyc^RNAi^* flies were significantly reduced in the sLN_v_s (Fig. 2B-C) and lLN_v_s (Fig. 2D-E). In contrast, PER levels were unaffected in the Dorsal Lateral Neurons (LN_d_s) (Fig. 2F-G). These results confirmed that, at least in a light-dark cycle (LD), *Pdf > cyc^RNAi^* flies have lower PER levels in *Pdf*^+^ neurons.

**Figure 2.**
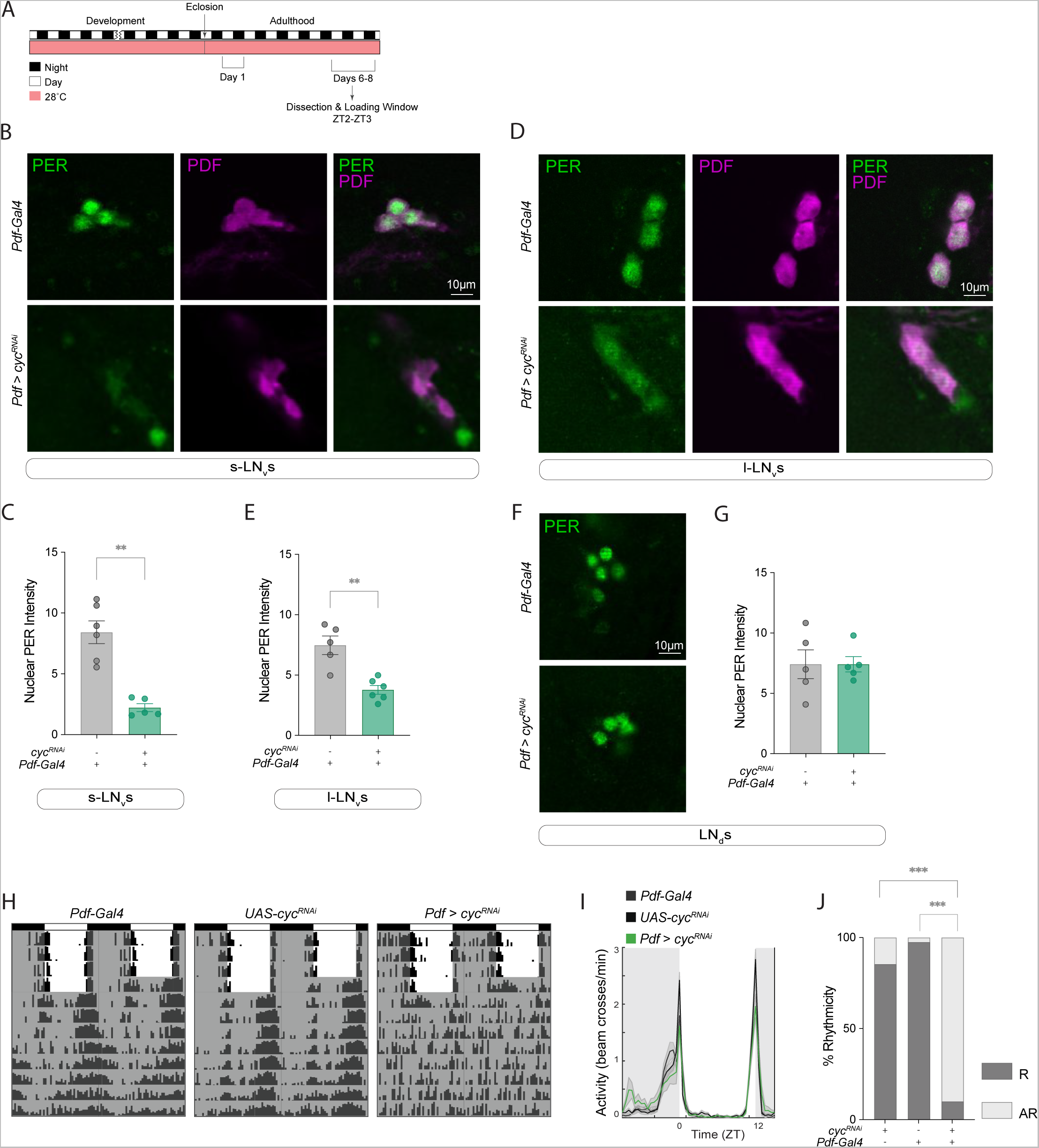
Constitutive *cyc* downregulation in *Pdf*+ cells leads to a reduction in PER levels and arrhythmicity under free-running conditions. (A) Representative timeline of the experiments in the figure. Flies were kept at 28°C for their entire lifespan. Experiments were then performed within Days 6-8 post-eclosion from pupae. Dissections were performed at ZT2-3. (B,D,F) Representative confocal images of PER (green) and PDF (magenta) staining in the sLN_v_s (B), lLN_v_s (D), and LN_d_s (F) of *Pdf > cyc^RNAi^*experimental and *Pdf-Gal4* /+ control flies (n = 5-6 brains per clock neuron group). All lines also included a *Pdf-RFP* transgene. Scale bar = 10 µm. (C,E,G) T-tests were used to compare nuclear PER intensity levels in the sLN_v_s (C), lLN_v_s (E), and LN_d_s (G) in flies of the indicated genotypes. Differences that are not significant are not indicated. ** p < 0.01. Error bars indicate SEM. (H) Representative actograms of flies of the indicated genotypes under 5 days of LD entrainment followed by 7 days of free-running (DD). To allow comparison with development-specific *cyc* downregulation, flies in this experiment were raised at 28°C for their entire lifespan and the experiment was conducted at 28°C. (I) Population activity plots for flies during days 3-5 of the LD cycle at 28°C. (J) Fisher’s exact contingency tests were used to analyze the percentage of rhythmic flies of the indicated genotypes under DD (DD1-7). The driver line also included a *tub-Gal80^ts^* transgene. Additional quantifications can be found in Table 1. R = Rhythmic and AR = arrhythmic. Differences that are not significant are not indicated. *** p < 0.001. Behavioral data correspond to two independent behavior experiments. For each genotype: 40 ≤ n ≤ 48.

**Table 1.** Summary of free running activity rhythms. Related to Figures 2, 3, and 7, and Supp. Figures 3, 4, and 6. Activity analysis of the above genotypes at either 28°C or 18°C. Light conditions for each experiment was 12:12 LD-DD. For each genotype the Number of Flies (n) is listed above. All flies were raised at 28°C or 18°C and then, upon eclosion, kept at 28°C or transferred to 18°C. ClockLab’s χ-square periodogram analysis was used to analyze rhythmicity, rhythmic power, and free-running period for each above genotypes. The % rhythmicity along with the number of rhythmic flies (nR), the period in hours with the SEM, and the rhythmic power with the SEM are indicated. Arrhythmic flies were not considered in the analysis.

| Temperature Pre-Ecdlosion: 28°C, Temperature Post Ecdlosion: 28°C |  |  |  |  |
| --- | --- | --- | --- | --- |
| Genotype | Number of Flies (n) | % Rhythmicity (nR) | Period (h) □ SEM | Rhythmic Power □ SEM |
| *;Pdf-Red,Pdf-Gal4; Tub-Gal80<sup>ts</sup>* | 40 | 97.50 (39) | 24.44 □ 0.06 | 109.80 □ 6.39 |
| *;;UAS-cyc<sup>RNAi 42563</sup>* | 48 | 85.42 (41) | 24.02 □ 0.08 | 85.74 □ 7.91 |
| *;Pdf-Red,Pdf-Gal4; Tub-Gal80<sup>ts</sup> > ;;UAS-cyc<sup>RNAi 42563</sup>* | 40 | 10.00 (4) | 25.88 □ 3.03 | 24.32 □ 3.71 |
| *;;UAS-Clk<sup>RNAi 42566</sup>* | 26 | 92.31 (24) | 23.73 □ 0.10 | 97.66 □ 8.90 |
| *;Pdf-Red,Pdf-Gal4; Tub-Gal80<sup>ts</sup> > ;;UAS-Clk<sup>RNAi 42566</sup>* | 21 | 52.38 (11) | 25.36 □ 0.22 | 36.74 □ 6.76 |
| *;Pdf-Red,Pdf-Gal4; Tub-Gal80<sup>ts</sup>* | 24 | 100.00 (24) | 24.5 □ 0.07 | 101.40 □ 8.97 |
| *;UAS-cas9/CyO; UAS-Vrig/TM6b Tb* | 16 | 87.50 (14) | 23.86 □ □ □ □ □ | □ □ □ □ □ □ □ □ □ |
| *;Pdf-Red,Pdf-Gal4; Tub-Gal80<sup>ts</sup> > ;UAS-cas9/CyO; UAS-Vrig/TM6b Tb* | 18 | 38.89 (7) | □ □ □ □ □ □ □ □ □ | 29.02 □ □ □ □ □ |
| *;Pdf-Red,Pdf-Gal4; Tub-Gal80<sup>ts</sup>* | 31 | 83.87 (26) | □ □ □ □ □ □ □ □ □ | 56.76 □ □ □ □ □ |
| *;UAS-Δcyc;* | 24 | 95.83 (23) | □ □ □ □ □ □ □ □ □ | 72.79 □ □ □ □ □ |
| *;Pdf-Red,Pdf-Gal4; Tub-Gal80<sup>ts</sup> > ;UAS-Δcyc;* | 32 | 3.13 (1) | □ □ □ □ □ □ □ □ □ | 13.66 □ □ □ □ □ |
| *;UAS-ΔClk* | 27 | 100.00 (27) | □ □ □ □ □ □ □ □ □ | 93.52 □ □ □ □ □ |
| *;Pdf-Red,Pdf-Gal4; Tub-Gal80<sup>ts</sup> > ;UAS-ΔClk* | 28 | 3.57 (1) | □ □ □ □ □ □ □ □ □ | 17.18 □ □ □ □ □ |
| Temperature Pre-Ecdlosion: 28°C, Temperature Post Ecdlosion: 18°C |  |  |  |  |
| Genotype | Number of Flies (n) | % Rhythmicity (nR) | Period (h) □ SEM | Rhythmic Power □ SEM |
| *;Pdf-Red,Pdf-Gal4 Tub-Gal80<sup>ts</sup>* | 45 | 84.44 (38) | 24.09 □ 0.09 | 100.7 □ 9.65 |
| *;;UAS-cyc<sup>RNAi 42563</sup>* | 62 | 96.77 (60) | 24.13 □ 0.05 | 80.98 □ 5.70 |
| *;Pdf-Red,Pdf-Gal4 Tub-Gal80<sup>ts</sup> > ;;UAS-cyc<sup>RNAi 42563</sup>* | 48 | 22.92 (11) | 23.64 □ 1.04 | 21.03 □ 2.66 |
| Temperature Pre-Ecdlosion: 18°C, Temperature Post Ecdlosion: 28°C |  |  |  |  |
| Genotype | Number of Flies (n) | % Rhythmicity (nR) | Period (h) □ SEM | Rhythmic Power □ SEM |
| *;Pdf-Red,Pdf-Gal4 Tub-Gal80<sup>ts</sup>* | 30 | 100 (30) | 24.93 □ 0.04 | 127.8 □ 6.76 |
| *;;UAS-cyc<sup>RNAi 42563</sup>* | 25 | 96 (24) | 24.04 □ 0.07 | 150.1 □ 12.44 |
| *;Pdf-Red,Pdf-Gal4 Tub-Gal80<sup>ts</sup> > ;;UAS-cyc<sup>RNAi 42563</sup>* | 31 | 38.71 (12) | 23.67 □ 0.61 | 24.27 □ 3.68 |

*cyc* null mutant flies have pronounced behavioral phenotypes. Their activity is unimodal instead of bimodal during LD, and they are predominantly nocturnal^47^. Additionally, *cyc* mutants are largely arrhythmic in DD due to the key role of *cyc* in circadian molecular oscillations^33^. We conducted behavioral experiments to determine the extent to which downregulating *cyc* specifically in PDF-expressing neurons recapitulates the phenotype of the *cyc* mutant. We found that at 28°C, the activity pattern of *Pdf > cyc^RNAi^* flies was still bimodal in LD (Fig. 2H-I). However, the majority (∼94%) of the experimental flies were arrhythmic in DD (Fig. 2J). *Pdf >* Δ*-cyc* flies showed similar behavioral phenotypes (Fig. S2 G-H), consistent with what was reported for their free-running behavior at 25°C^43^.

### *cyc* acts during development to shape neuronal morphology in adults

To knock down *cyc* specifically during development, we employed a temperature-sensitive Gal80 (Gal80^ts^) variant with ubiquitous expression to conditionally inhibit Gal4-mediated expression of the RNAi^48^. This method enables the temporal regulation of UAS transgenes, as Gal80^ts^ remains active at lower temperatures but becomes inactive at higher temperatures. We raised flies at 28°C to allow *cyc* downregulation during development then transferred them to 18°C immediately after eclosion (Fig. 3A). After 1 week at 18°C, the brains were dissected at ZT2 and stained with PDF and RFP antibodies (see methods section). As shown in Figure 3, downregulating *cyc* exclusively before eclosion resulted in abnormal morphology of the sLN_v_ axonal projections in adult flies (Fig. 3B). The phenotypes resembled those observed with constitutive downregulation, with a significantly shorter distance to the branching point (Fig. 3B-C) and a greater degree of defasciculation compared to parental controls (Fig. 3B-D). No significant differences were found in the total projection length or the degree of defasciculation of the lLN_v_s (Fig. 3E-F).

**Figure 3.**
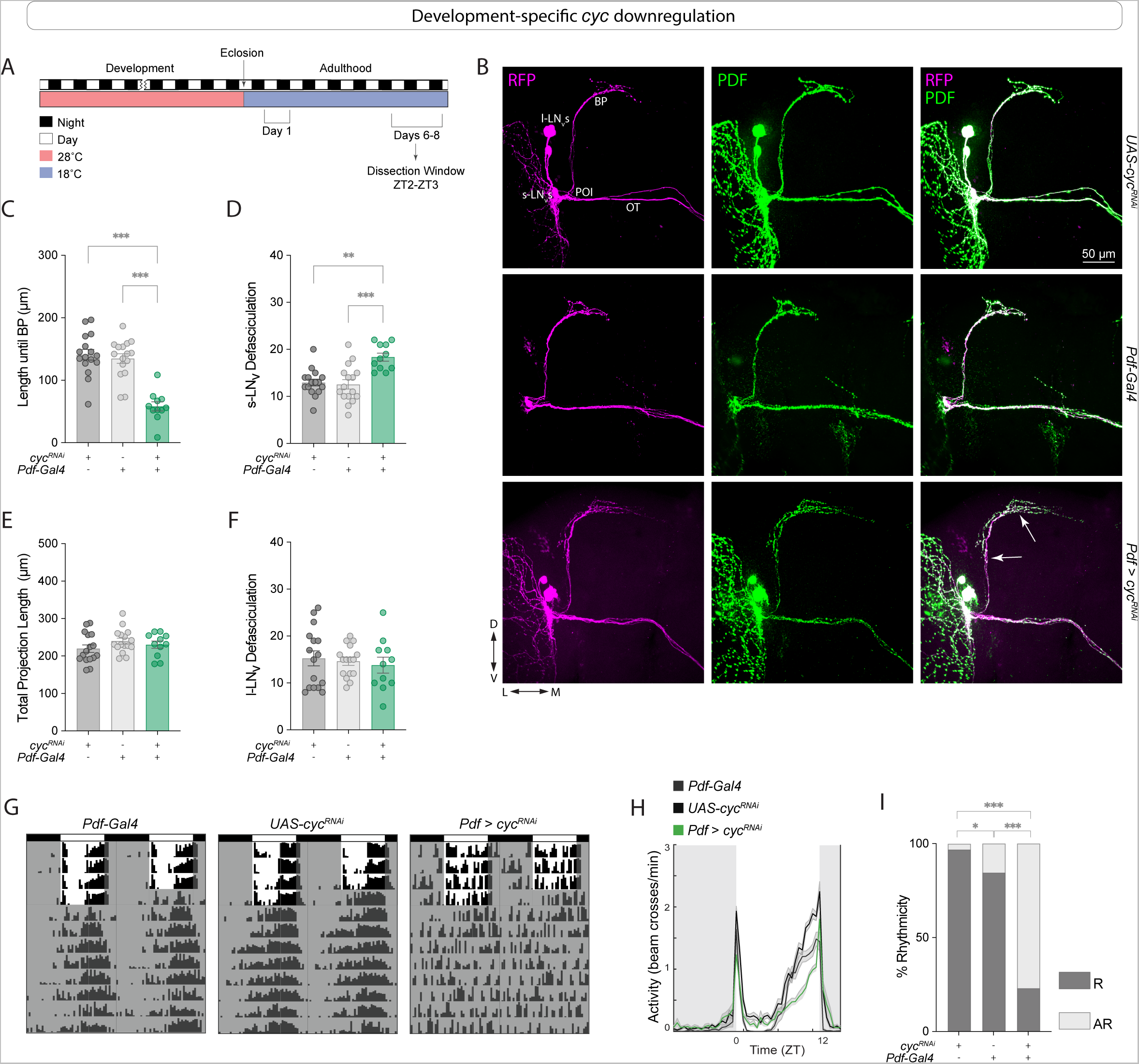
Development-specific *cyc* downregulation in *Pdf*+ cells prevents sLN_v_ fasciculation. (A) Representative timeline of the experiments in the figure. Flies were raised in LD at 28 °C, and transferred to 18 °C immediately after eclosion. Dissections were then performed within Days 6-8 post-eclosion from pupae at ZT2-3. (B) Representative confocal images of anti-PDF (green) and anti-RFP (magenta) staining of adult fly brains in which *cyc* was downregulated only during development. Each line also included a *Pdf-RFP* transgene. The sLN_v_s, lLN_v_s, OT, BP, and POI are labeled. Scale bar = 50 µm. (C-F) Quantification of the LN_v_ morphology phenotypes of flies of the indicated genotypes. The driver line also included a *tub-Gal80^ts^*transgene. The sLN_v_ projection length until the branching point (BP) (C), the number of intersections of sLN_v_ ventral projections (D), the total sLN_v_ projection length (E), and the total number of intersections of the lLNv projections along the optic tract (OT) (F) are shown for flies in which *cyc* was downregulated in *Pdf*+ cells until eclosion. Two independent experiments were conducted. For each genotype: 11 ≤ n ≤ 16. Kruskal-Wallis tests followed by Dunn’s multiple comparisons tests were used to quantify the LN_v_ morphology. ** p < 0.01, *** p < 0.001. Error bars indicate SEM. Each dot corresponds to one brain. (G-I). Behavioral phenotypes of development-specific *cyc* knockdown. Flies were raised in LD at 28 °C, before being transferred to 18°C upon eclosion. Experiments were conducted at 18°C. (G) Representative actograms of flies of the indicated genotypes under free-running (see Table 1 for n and additional quantifications). (H) Population activity plots for flies during days 3-5 of the LD cycle at 18°C. (I) Percent rhythmicity for the indicated genotypes under DD. R=Rhythmic and AR= arrhythmic. Fisher’s exact contingency tests were used to analyze the percentage of rhythmic flies under DD (DD1-7). * p < 0.05, *** p < 0.001. Error bars indicate SEM. The data correspond to two independent behavior experiments. For each genotype: 45 ≤ n ≤ 62.

A previous study showed that panneuronal rescue of *cyc* expression in a *cyc^01^* mutant exclusively during development was sufficient to partially rescue arrhythmicity in adult flies^40^. Therefore, we asked if downregulating *cyc* in the *Pdf*+ cells specifically during development would lead to behavioral phenotypes similar to those seen in the *cyc* null mutants. We found that under free-running conditions at 18°C, most (∼78%) of the *Pdf > cyc^RNAi^* flies were arrhythmic (Fig. 3G-I). These results indicate that developmental downregulation of *cyc* specifically in the *Pdf*+ cells is sufficient to prevent behavioral rhythms in adults.

To determine whether adult-specific *cyc* knockdown in the *Pdf*-expressing cells would also lead to morphological phenotypes, we raised flies at 18°C and switched them to 28°C immediately after eclosion (Fig. S3A, see the methods section). This manipulation did not result in morphological phenotypes either in terms of the length to the branching point or in the degree of sLN_v_ defasciculation (Fig. S3B-F). Under free-running 28°C, the majority of the experimental flies were arrhythmic (Fig. S3 G-H), indicating that, as expected, *cyc* is required in adult clock neurons for proper circadian clock function.

### *Cyc* manipulations lead to aberrant sLN_v_ projections in larval clock neurons

Next, we asked if *cyc* downregulation results in clock neuron morphology phenotypes during earlier developmental stages. The four larval sLNvs, which modulate the sensitivity of larvae to light and mediate a circadian rhythm in visual sensitivity^49^, appear to be identical in their anatomy and synaptic connections^50^. We expressed the *cyc*^RNAi^ transgene under the *Pdf-Gal4* driver and dissected third larval instar (L3) brains (Fig. 4). In brains of experimental larvae the length to the branching point did not differ from that of the controls (Fig. 4B-C), but the degree of dorsal termini branching was significantly higher (Fig. 4B-D). This quantification is similar to that previously described when quantifying the arborization of the dorsal projections sLN_v_s in adults^27^, where the concentric circles are centered at the main dorsal branching point (Fig. S1B; see methods section). The total sLN_v_ projection length was not affected by the genetic manipulation (Fig. 4E).

**Figure 4.**
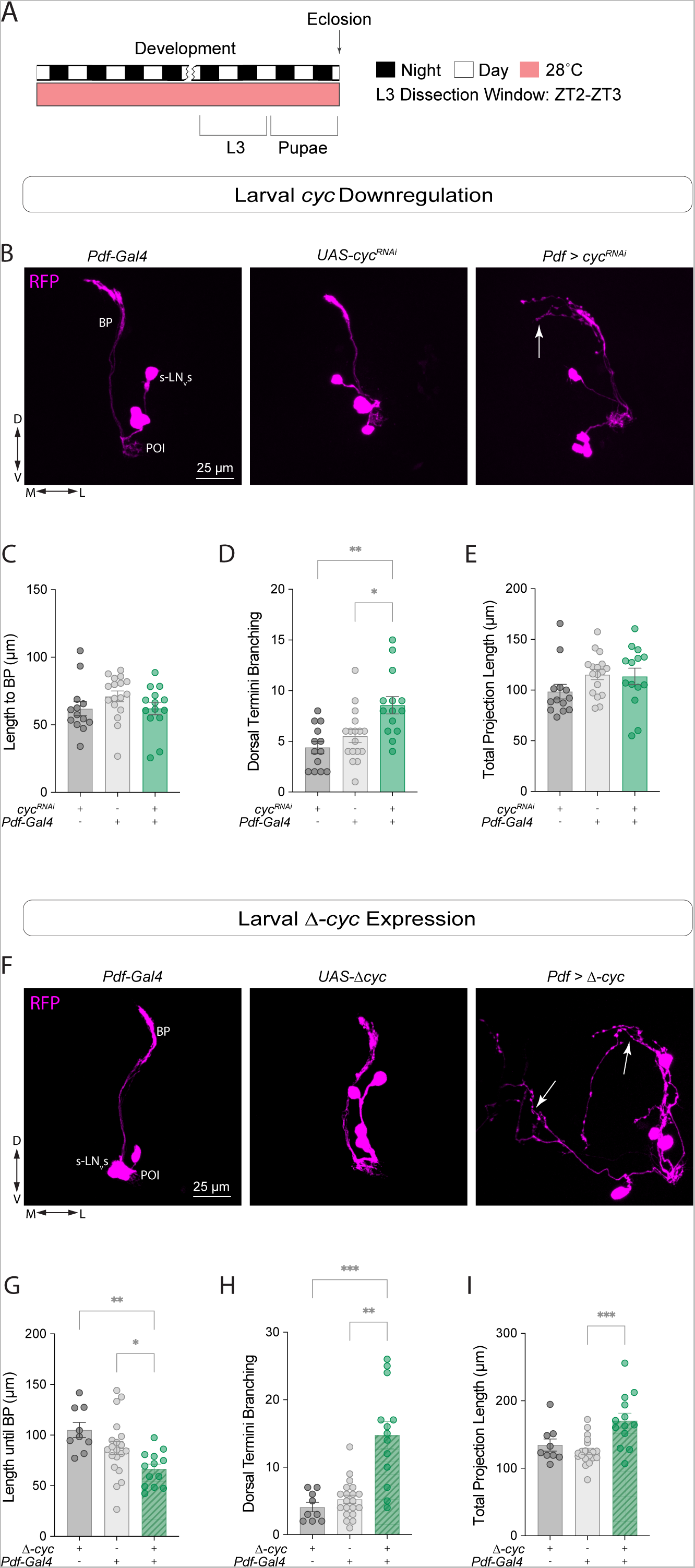
*Cyc* manipulations lead to aberrant sLN_v_ projections in larval clock neurons. (A) Representative timeline of the experiments in the figure. Larvae were raised in LD at 28 °C. Third instar larvae (L3) were dissected at ZT2-3. (B-E) Developmental effects of *cyc* knockdown in the sLN_v_s. (B) Representative confocal images of L3 larval brains stained with anti-RFP (magenta) labeling the sLN_v_s. The sLN_v_s, POI, and BP are labeled. Scale bar = 25 µm. (C-E). Kruskal-Wallis tests followed by Dunn’s multiple comparisons tests were used to compare the projection length from the POI to the BP (C), the degree of sLN_v_ dorsal termini branching (D), and the total projection length (E). * p < 0.05, ** p < 0.01. Each dot corresponds to one brain. For each genotype: 13 ≤ n ≤ 17. (F-I) Developmental effects of expressing a dominant-negative form of *cyc*, Δ*-cyc*, in the larval sLN_v_s. (F) Representative confocal images of anti-RFP (magenta) staining in the sLN_v_s of L3 larvae. Kruskal-Wallis tests followed by Dunn’s multiple comparisons tests compared the projection length from the POI to the BP (G), the degree of sLN_v_ dorsal termini branching (H) and the total projection length (I). Each dot corresponds to one brain. For each genotype: 9 ≤ n ≤ 20. * p < 0.05, ** p < 0.01, *** p < 0.001. Three independent experiments were conducted for each genetic manipulation and each line also included a *Pdf-RFP* transgene. The driver lines also included a *tub-Gal80^ts^* transgene. Error bars indicate SEM.

Since the effects of *cyc* knockdown via RNAi and the expression of a *cyc* dominant negative form in adults were similar (Fig. 1 and Fig. S2), we analyzed the morphology of the sLN_v_ projections in L3 larvae upon Δ*-cyc* expression. In *Pdf >* Δ*-cyc* larval brains, the length to the branching point was significantly lower (Fig. 4F-G) and the number of branches was significantly greater than that of controls (Fig. 4H). The total projection length was not affected (Fig. 4I). Taken together, these results suggest that *cyc* plays a role in the development of the larval sLN_v_ neurons.

### *Clk* and *cyc* have differential effects on neuronal morphology

*Clk* and *cyc* mutations produce similar effects on the expression pattern of PDF in adult brains^23^. CLK and CYC act as heterodimeric transcriptional activators, and the circadian phenotypes associated with mutations in these core circadian clock genes, both molecular and behavioral, are largely similar^43,47,51^. To determine if downregulating *Clk* in the sLN_v_s leads to the same defasciculation of the sLN_v_s observed with *cyc* manipulations, we performed similar experiments as those described above, in which we expressed *Clk*^RNAi^ in *Pdf*+ neurons. We found that *Pdf > Clk^RNAi^* flies also showed neuronal morphology phenotypes (Fig. 5).

**Figure 5.**
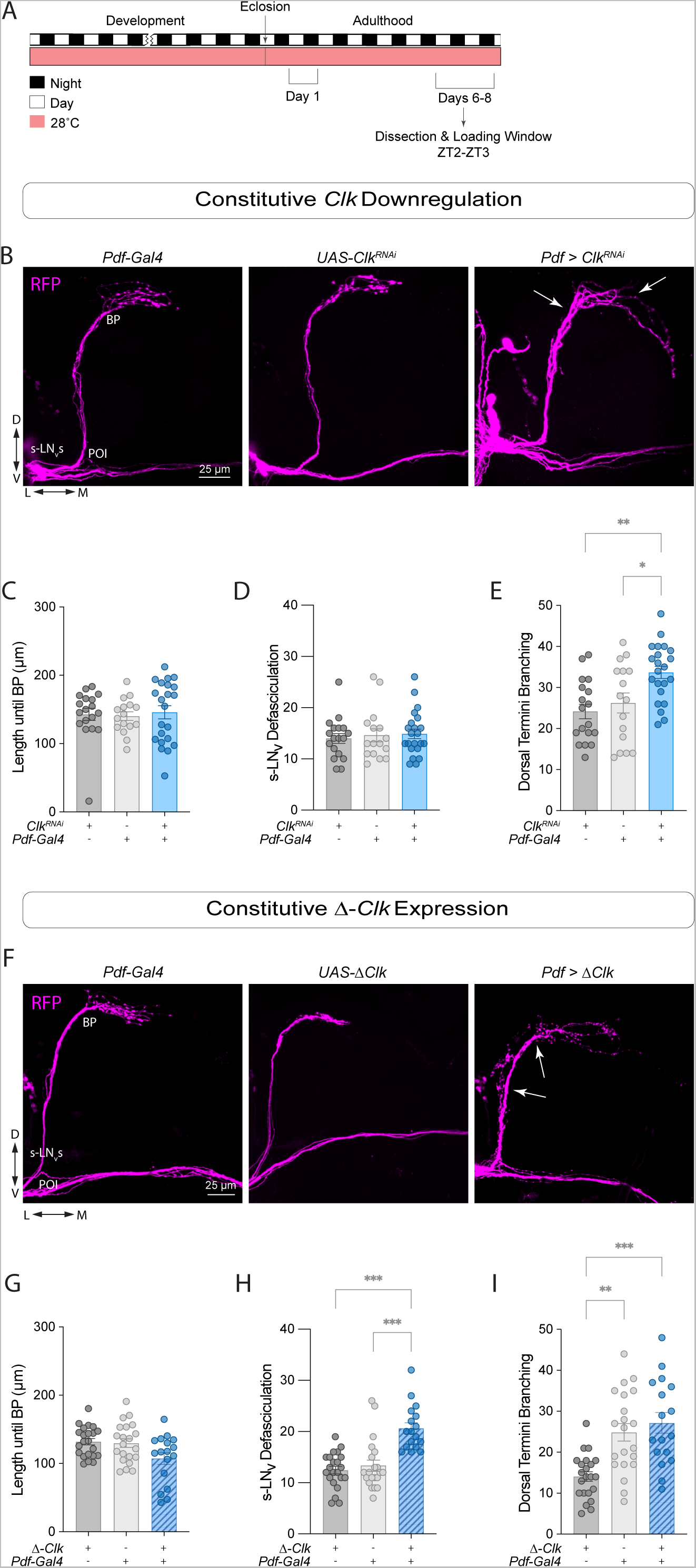
*Clk* and *cyc* manipulations result in different morphology phenotypes in clock neurons. (A) Representative timeline of the experiments in the figure. Flies were kept in LD conditions at 28 °C for their entire lifespan. Dissections were performed within Days 6-8 post-eclosion from pupae at ZT2-3. (B) Representative confocal images of anti-RFP (magenta) staining in the sLN_v_s adult brains of control (*Clk^RNAi^ /*+ and *Pdf-Gal4;tub-Gal80^ts^/*+), and experimental (*Pdf* > *Clk^RNAi^*) flies. All lines employed also included a *Pdf-RFP* transgene. Scale bar = 25 µm. (C-E) Quantification of sLN_v_ morphology using Kruskal-Wallis tests followed by Dunn’s multiple comparisons tests, compared the length until the branching point (C), the total number of axonal crosses of the sLN_v_s (D), and the total number of axonal crosses after the BP (E). Each dot corresponds to one brain. Two independent experiments were conducted. For each genotype: 16 ≤ n ≤ 22. * p < 0.05, *** p < 0.001. Error bars indicate SEM. (F) Representative confocal images of anti-RFP (magenta) staining in the sLN_v_s adult brains of control (*UAS-*Δ*Clk/*+ and *Pdf-Gal4;tub-Gal80^ts^/*+), and experimental (*Pdf* > Δ*-Clk*) flies. All lines employed also included a *Pdf-RFP* transgene. Scale bar = 25 µm. (G-I) Quantification of sLN_v_ morphology using Kruskal-Wallis tests followed by Dunn’s multiple comparisons tests, compared the length until the branching point (G), the total number of axonal crosses of the sLN_v_s (H), and the total number of axonal crosses after the BP (I). Each dot corresponds to one brain. Two independent experiments were conducted. For each genotype: 17 ≤ n ≤ 22. ** p < 0.01, *** p < 0.001. Error bars indicate SEM.

In a previous study, *Clk* downregulation resulted in overfasciculation of the sLN_v_ dorsal termini when stained with anti-PDF^52^. However, RFP labeling of the sLN_v_ membrane indicated that these termini were actually more expanded than those of control flies, resulting in significantly higher dorsal termini branching (Fig. 5B-E and Supl.Fig. 1C). In *Pdf > Clk^RNAi^* flies, neither the distance to the branching point (Fig. 5C) nor the degree of defasciculation differed from controls (Fig. 5D). Neither the sLN_v_ total projection length nor the lLN_v_ projections were affected (Fig. S4C-D). Only ∼48% of the *Pdf > Clk^RNAi^* flies were rhythmic, and those that were rhythmic exhibited a lengthening of the free-running period (Fig. S4B, Table 1). Expression of Δ*-Clk* in the *Pdf*+ cells did not result in changes in the sLN_v_ projection length until branching point (Fig. 5F-G) or the total length of the projections (Fig. S4F). However, the *Pdf* > Δ*-Clk* brains had increased defasciculation of the ventral projection (Fig. 5H). The degree of dorsal termini branching in the *Pdf* > Δ*-Clk* flies was not significant (Fig. 5I), not was the degreed of lLN_v_ defasciculation (Fig. S4G). Under DD at 28°C, the majority of Δ*-Clk* expressing flies were arrhythmic (Fig. S4E), consistent with what was reported at 25°C^43^.

We then examined L3 larval brains to determine if the observed phenotypes were already present at this developmental stage. While expression of *Clk^RNAi^* did not result in morphological phenotypes in larval LN_v_s (Fig. S5), expression of Δ*-Clk* resulted in pronounced phenotypes (Fig. 6A-B). We observed a significant increase in sLN_v_ dorsal termini branching (Fig. 6D) and total projection length (Fig. 6E) in *Pdf >* Δ*-cyc* larvae. The length to the branching point for the experimental larvae was not significantly different from that of the control lines (Fig. 6C). Expressing Δ*-Clk* led to more pronounced phenotypes in the larval stage than *Clk* downregulation, possibly due to incomplete knockdown.

**Figure 6.**
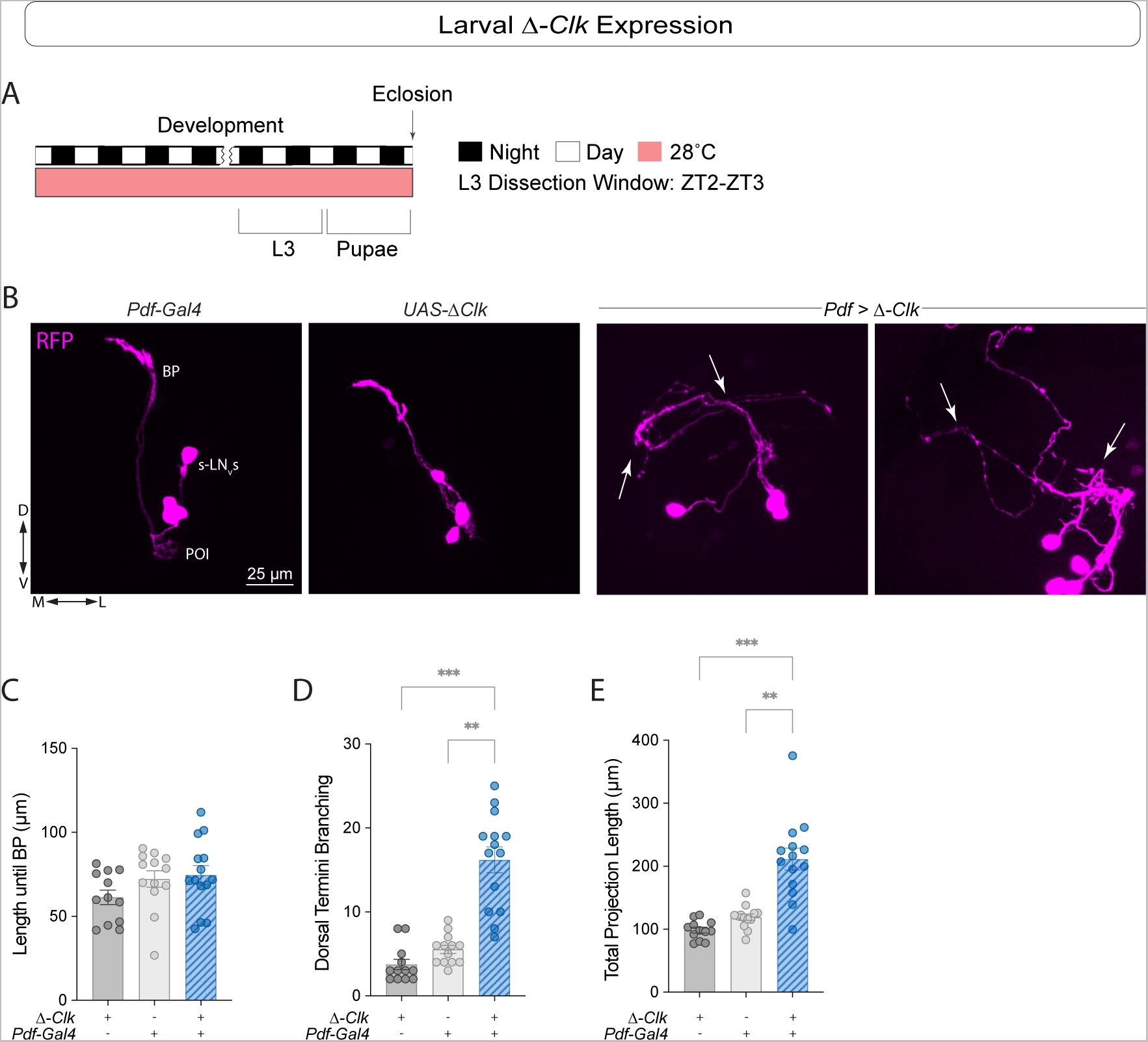
Expressing Δ*-Clk* in the sLN_v_s leads to axonal morphology phenotypes in L3 larvae. (A) Representative timeline of the experiments in the figure. Larvae were raised in LD at 28 °C. Third instar larvae (L3) were dissected at ZT2-3. (B) Representative confocal images of anti-RFP (magenta) staining in the sLN_v_s when Δ*-Clk* was expressed in *Pdf*+ neurons in L3 larvae. Each line also included a *Pdf-RFP* transgene and the driver line also included a *tub-Gal80^ts^* transgene. Scale bar = 25 µm. Kruskal-Wallis tests followed by Dunn’s multiple comparisons tests were used to compare the length to the BP (C), the degree of sLN_v_ dorsal termini branching (D), and the total projection length (E). Two independent experiments were conducted. Each dot corresponds to one brain. For each genotype: 12 ≤ n ≤ 14. * p < 0.05, ** p < 0.01, *** p < 0.001. Error bars indicate SEM.

In addition to the main feedback loop, CLK and CYC form a secondary loop by activating *vri* and *Pdp1ε*^8,9^, which repress and activate *Clk* expression, respectively. The low PDF peptide in the sLN_v_s projections of *cyc^01^* mutants can be rescued by *vri* overexpression^39^. To determine if *vri* expression also affects sLN_v_s morphology we expressed a line with a CRISPR/Cas9-based gRNA targeting the *vri* gene^53^ under the control of the *Pdf*-Gal4 driver. We found that in *Pdf* > Cas9 + *vri-g* flies neither the distance until branching point in the s-LN_v_s (Fig. 7B-C) nor the degree of fasciculation of the s-LN_v_s (Fig. 7D) was different from controls. The total projection length was significantly higher than controls (Fig. 7E), but in this case due to projections extending ventrally towards the optic tract (Fig. 7B). In the majority of the brains, some s-LN_v_ projections extended towards the ventral brain after reaching the SMP (Fig. 7B) and in most cases contacted the l-LNv contralateral projections in the optic tract (Fig. 7F-G), a phenotype that was never observed in control brains.

**Figure 7.**
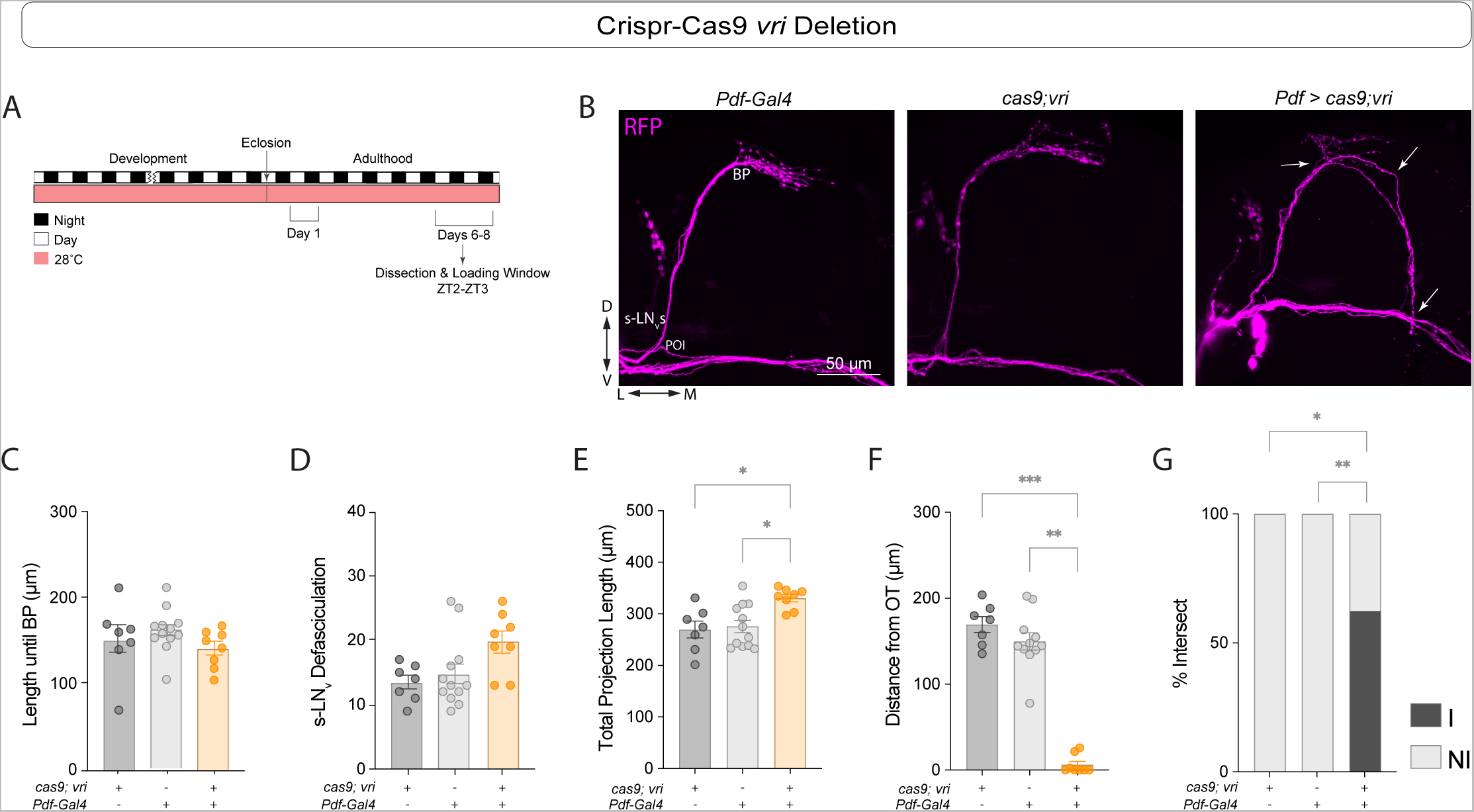
*Vri* mutagenesis results in sLN_v_ hyperextension. (A) Representative timeline of the experiments in the figure. Flies were kept in LD conditions at 28 °C for their entire lifespan. Dissections were performed within Days 6-8 post-eclosion from pupae at ZT2-3. Behavioral experiments were run at 28°C. (B) Representative confocal images of anti-RFP (magenta) staining in the sLN_v_s adult brains of control (*cas9;vrig\/*+ and *Pdf-Gal4;tub-Gal80^ts^/*+), and experimental (*Pdf* > *cas9;vrig*) flies. All lines employed also included a *Pdf-RFP* transgene. Scale bar = 25 µm. (C-E,G,H) Kruskal-Wallis tests followed by Dunn’s multiple comparisons tests compared the length until the branching point (C), the total number of axonal crosses of the sLN_v_s (D), the longest path of the sLN_v_ neurites (without including misrouting) (E), and the distance from the sLN_v_ projection end to the lLN_v_s projection along the optic tract (F). (H) Fisher’s exact contingency tests were used to analyze the percentage of brains where the sLN_v_s intersected with the lLN_v_s in the optic tract (I = Intersecting, N.I. = Not Intersecting). Each dot corresponds to one brain. Two independent experiments were conducted. For each genotype: 7 ≤ n ≤ 12. * p < 0.05, ** p < 0.01, *** p < 0.001. Error bars indicate SEM.

## Discussion

Our results reveal a role for the circadian clock gene *cyc* in establishing the proper cellular morphology of the key clock pacemaker neurons, the sLN_v_s. Both constitutive *cyc* knockdown or expression of a dominant negative form of *cyc* in *Pdf*+ cells result in increased defasciculation of the sLN_v_s. In addition, *Clk* downregulation and expression of a dominant negative form of *Clk* also result in sLN_v_ morphology phenotypes, although some of those phenotypes appear to be distinct from those caused by *cyc* manipulations. Expressing the dominant-negative forms of either *Clk* or *cyc* has been used in previous studies as an effective way to prevent molecular oscillations in subsets of clock neurons. However, our results indicate that these genetic manipulations lead to additional morphological phenotypes beyond molecular timekeeping that are already detectable during the larval stages.

In addition to anatomical and functional classifications, clock neurons can be divided into early or late developmental groups depending on when circadian oscillations can be detected. In the early groups, which include the sLN_v_s, *per* and *tim* expression rhythms can be detected at the first instar (L1) larval stage, whereas in the late groups, such rhythms cannot be detected until metamorphosis^25,54^. However, *cyc* and *Clk* expression using GFP-*cyc* and GFP-*Clk* transgenes can be detected in almost all groups of clock neurons at early developmental stages, even days before *per* oscillations begin ^10^. This suggests that *cyc* and *Clk* play additional roles in the development of clock neurons beyond their role in the molecular oscillator.

Both *cyc* and *Clk* modulate PDF expression in both larval and adult clock neurons. In *Clk*^jrk^ mutants, neither PDF nor *pdf* mRNA can be detected in larval^32^ or adult LN_v_s^23^, and similar effects have been observed for the *cyc^02^* mutant^23^. A previous study showed that panneuronal rescue of *cyc* expression throughout development is sufficient to restore PDF expression in the LN_v_ dorsal projections of *cyc^01^* mutants and that the developmental inhibition of CLK/CYC activity affects PER expression rhythms in adult LN_v_s^40^. Moreover, overexpression of *vri*, a clock gene that is downstream of CLK/CYC and acts as a repressor of CLK transcription^9,32^, causes a severe reduction in PDF levels in larval brains^32^, and the low PDF levels in the sLN_v_s of *cyc^01^* mutants can be rescued by *vri* overexpression^39^. However, restoring PDF expression in the sLN_v_s in flies lacking *vri* expression is not sufficient to rescue activity rhythms.

Unlike in *cyc* and *Clk* mutants^23^, PDF can be detected in the sLN_v_s projections in *per^01^* and *tim^01^* mutants, although it no longer shows rhythms in its accumulation in the dorsal termini^23^. In addition, structural plasticity rhythms in the sLN_v_s are absent in both *per^01^* and *tim^01^* mutants^27^. Downregulation of *Clk* ^52^, expression of Δ*-cyc* ^55^, and overexpression of *vri* in *Pdf*-expressing cells^39^ also result in impaired plasticity rhythms^56^. Although the anatomical phenotypes seen in these mutants are milder than those that observed when *cyc* and *Clk* are downregulated or when their dominant negative forms are expressed, the sLN_v_ projections of both *per* and *tim* null mutants also exhibit altered morphology^27^.

Expression of CLK and CYC proteins is restricted to clock neurons^10^; however, *cyc* mRNA is more widely expressed^57^. Expressing *Clk* outside the clock network leads to the generation of ectopic clocks^58^, but these require *cyc*, suggesting that although the CYC protein can be detected only in clock cells, *cyc* mRNA is more broadly expressed. A study by Liu et al. showed that *Clk* stabilizes CYC both in cultured *Drosophila* Schneider 2 (S2) cells and *in vivo*: upon ectopic *Clk* expression, GFP-CYC can be detected in additional cells beyond the clock neuron network^59^. The role of *cyc* mRNA expression in non-clock cells remains unknown. Our results suggest that *Clk* and *cyc* are involved in shaping the morphology of clock neurons, and it is possible that they play similar roles in non-clock neurons as well.

## Materials and Methods

### Fly lines and rearing

Flies were raised on cornmeal-sucrose yeast media in a Percival Incubator under 12:12 LD at different temperature conditions. Depending on the experiment, flies were raised under either 18°C, 25°C, or 28°C (indicated in the figure legends). The lines *UAS-cyc^RNAi^* (BDSC #42563)*, UAS-Clk^RNAi^* (BDSC #42566), *w*^1118^ (BDSC #3605) and CS (BDSC #64349) were obtained from the Bloomington *Drosophila* Stock Center. The lines *Pdf-RFP,Pdf-Gal4;Tub-gal80^ts^* and *w;Pdf-RFP;MKRS/TM6* were donated by Justin Blau (New York University). The *cyc^01^*, UAS-Δ*-cyc*, and UAS-Δ*-Clk* stocks were donated by Paul Hardin (University of Texas).

### Immunohistochemistry

#### LN_v_ PDF levels and neuronal morphology

Brains of 6–8-day-old adult males or L3 larvae were dissected between ZT2 and ZT3 in ice-cold Schneider’s *Drosophila* Medium (S2) (Thermo Fisher, #21720024). They were fixed immediately after dissection in 2% Paraformaldehyde (PFA) in S2 for 30 minutes. Brains were then treated with blocking solution (5% goat serum in 0.3% PBS-Tx) for 1 hr at room temperature followed by incubation with primary antibodies at 4°C for 24–48 hr. The primary antibodies used were 1:3000 mouse anti-PDF (Developmental Hybridoma Bank) and 1:1000 rabbit anti-RFP (Rockland, #600-401-379-RTU). After incubation, the brains were rinsed 6 times in 0.3% PBS + Triton X-100 (PBT), after which they were incubated with Alexa-fluor conjugated secondary antibodies for 24-hr at 4°C. The secondary antibodies used were 1:3000 Alexa-488 (Thermo Fisher, #A11029) and 1:1000 Alexa-568 (Thermo Fisher, #A11036). The brain samples were further washed 6 times with 0.3% PBT, cleaned and mounted on a clean glass slide in Vectashield (Vector Laboratories, #H-1000-10) mounting media.

#### PER Staining

Brains of 6–8-day-old males were dissected one hour before lights-on (ZT23) in ice-cold Schneider’s *Drosophila* Medium (S2) (Thermo Fisher, #21720024). Immediately after dissection, brains were fixed in 2% paraformaldehyde (PFA) for 30 minutes, stained and mounted as described above. The primary antibodies used were 1:1000 rabbit anti-RFP (Rockland, #600-401-379-RTU) and 1:500 rat anti-PER (donated by Orie Shafer). The secondary antibodies used were 1:1000 Alexa-568 (Thermo Fisher, #A11036) and 1:500 Alexa-488 (Thermo Fisher, #A21208).

### Imaging, quantification, and Statistical Analysis

All images were acquired on an Olympus Fluoview 1000 laser-scanning confocal microscope using a 40x/1.10 NA FUMFL N objective (Olympus, Center Valley, PA) at the Advanced Science Research Center (ASRC-CUNY). For all the experiments, only one hemisphere per brain was imaged (the right hemisphere, unless it was damaged, in which case we imaged the left hemisphere).

#### Quantification of adult LN_v_ morphology

We quantified the total projection length of the sLN_v_s, the axonal projection length from the point of origin (POI) until the branching point, the degree of defasciculation of the sLN_v_ ventral projections (Fig. S1A), and the dorsal termini branching (Fig. S1C). If the axonal projection length went past the midline of the brain, the length was measured up to the midline. The projection length and partial projection length of the sLN_v_s were quantified using Fiji [31] in ImageJ [32]. The length of the dorsal projection was determined by a line drawn from the point of intersection (POI) between the sLN_v_s and the optic tract until the end of the dorsal termini. The partial length of the dorsal projections was determined by a line drawn from the same point of intersection until the branching point (BP) of the sLN_v_s at the dorsal termini. A modified Scholl’s analysis [21] was used to measure the defasciculation of the sLN_v_ projections and the optic tract projection, which corresponds to the lLN_v_s. (Fig. S1A and 1C, respectively). Six concentric circles, each 25 microns (µm) apart, were placed around the same point of intersection used in the length measurements.

#### Quantification of larval sLN_v_ morphology

We quantified the total projection length of the sLN_v_s, the axonal projection length until the branching point of the and the degree of branching in the sLN_v_ dorsal projections. The projection length, partial projection length, and area of the sLN_v_s were quantified using the Fiji platform [31] in ImageJ [32]. The projection length was measured by a line drawn from a determined first point of intersection (POI) of each of the sLN_v_ cell bodies until the end of the dorsal termini. The partial length of the axonal projections was determined by a line drawn for the same point of intersection until the branching point (BP) of the sLN_v_s at the dorsal termini. A modified Scholl’s analysis was used to measure the branching of the sLN_v_ projections. Six concentric circles were placed around the same branching point used in the length measurements. The concentric circles were each 12.5 µm away from each other, so that the farthest circle was 75 µm away from the POI. The number of visible neurites of the sLN_v_s that intersected with each circle were counted and summed, yielding the total number of intersecting neurons for the dorsal projections.

#### Quantification of PER levels

Single optical sections of either sLN_v_s, lLN_v_s or LN_d_s were imaged using the same settings using a 40x/1.10 objective. PDF was used to identify the small and large LN_v_s. The LN_d_s were identified based on their localization, size, and morphology. PER levels were determined through normalization of nuclear staining within each cell to the background. The average value for each brain within a cluster was computed by averaging the values obtained from multiple cells within that cluster. Quantification was performed using images from 5-6 brains per each cluster at each timepoint.

### Locomotor activity rhythm recording and analysis

DAM2 *Drosophila* Activity Monitors (TriKinetics, Waltham, MA) were used to record the locomotor activity rhythms of adult male flies aged three-to five-days, as previously described ^60^. Flies were entrained to 12:12 LD cycles for at least five days, and then transferred to constant darkness (DD) for at least eight days at a constant temperature of 28°C, unless otherwise specified. Free-running activity rhythms were analyzed with ClockLab software from Actimetrics (Wilmette, IL). We employed ClockLab’s χ-square periodogram function, which was integrated into ClockLab software, for the analysis of rhythmicity, rhythmic power, and free-running period in individual flies, using a confidence level of 0.01 [33]. For each of the tested genotypes, only significant periodicities falling within the 14 to 34-hour range were taken into consideration. In instances where an individual fly exhibited multiple periodicities with peaks surpassing the significance threshold, only the period with the highest amplitude was utilized when calculating the average periods presented in Table 1. ClockLab assigns each peak in the χ-square periodogram both a “Power” value and a “Significance” value. The “Rhythmic Power” for each designated rhythmic fly was determined by subtracting the “Significance” value from the “Power” value associated with the predominant peak. Flies that did not exhibit a periodicity peak above the threshold (10) were categorized as “arrhythmic,” and their period and rhythmic power were not included in the analysis ^60^.

### Statistical Analysis

Pearson’s D’Agostino normality tests were performed for all the datasets. Depending on whether the data were normally distributed, statistical analyses were performed using either a one-way ANOVA with a Tukey’s multiple comparisons test or a Kruskal-Wallis test with a Dunn’s multiple comparisons test for 3 or more groups, or a t-test for comparisons between 2 groups. Fisher’s exact contingency tests were run to analyze the percent rhythmicity for the indicated genotypes under DD.

## Materials and Reagents

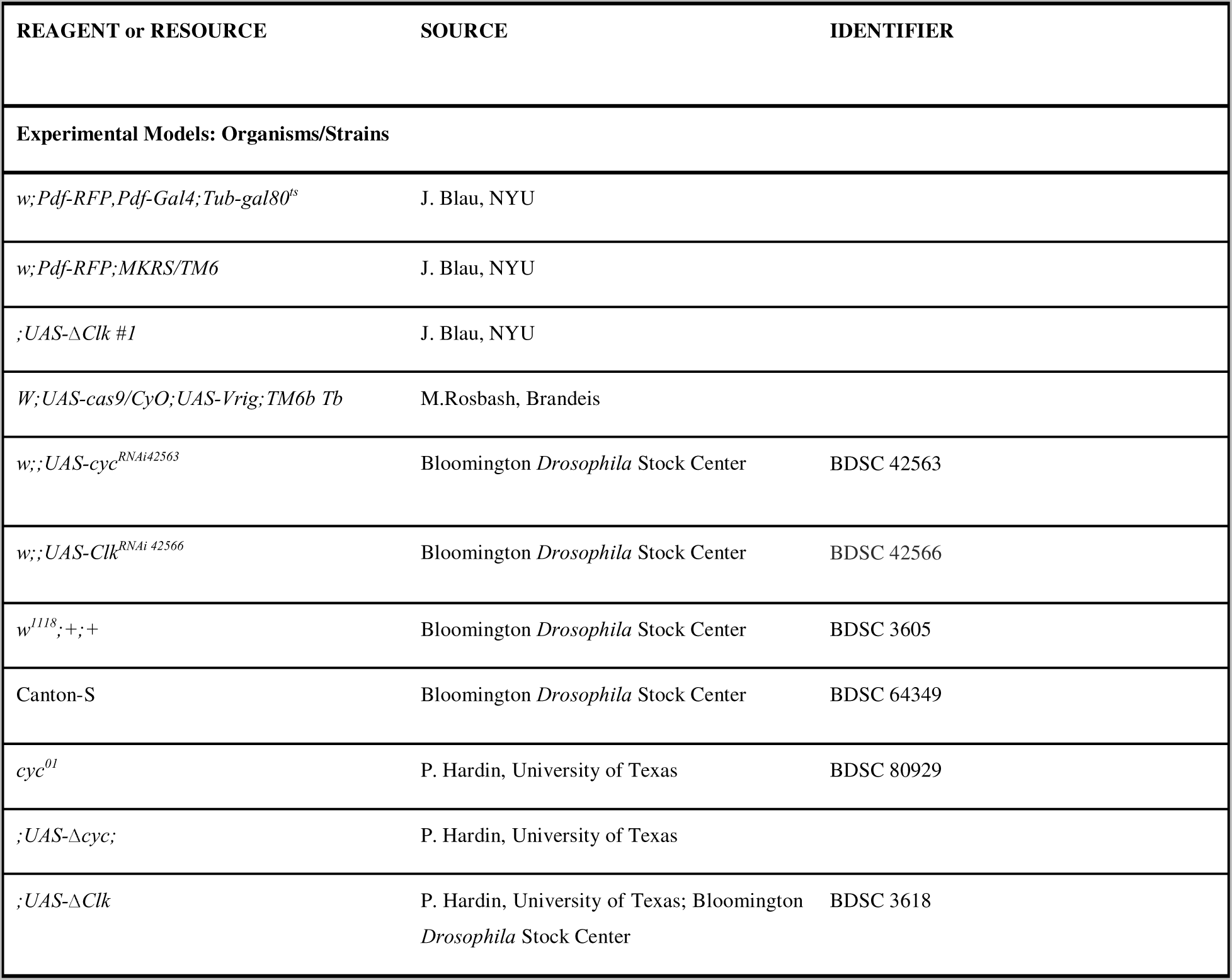
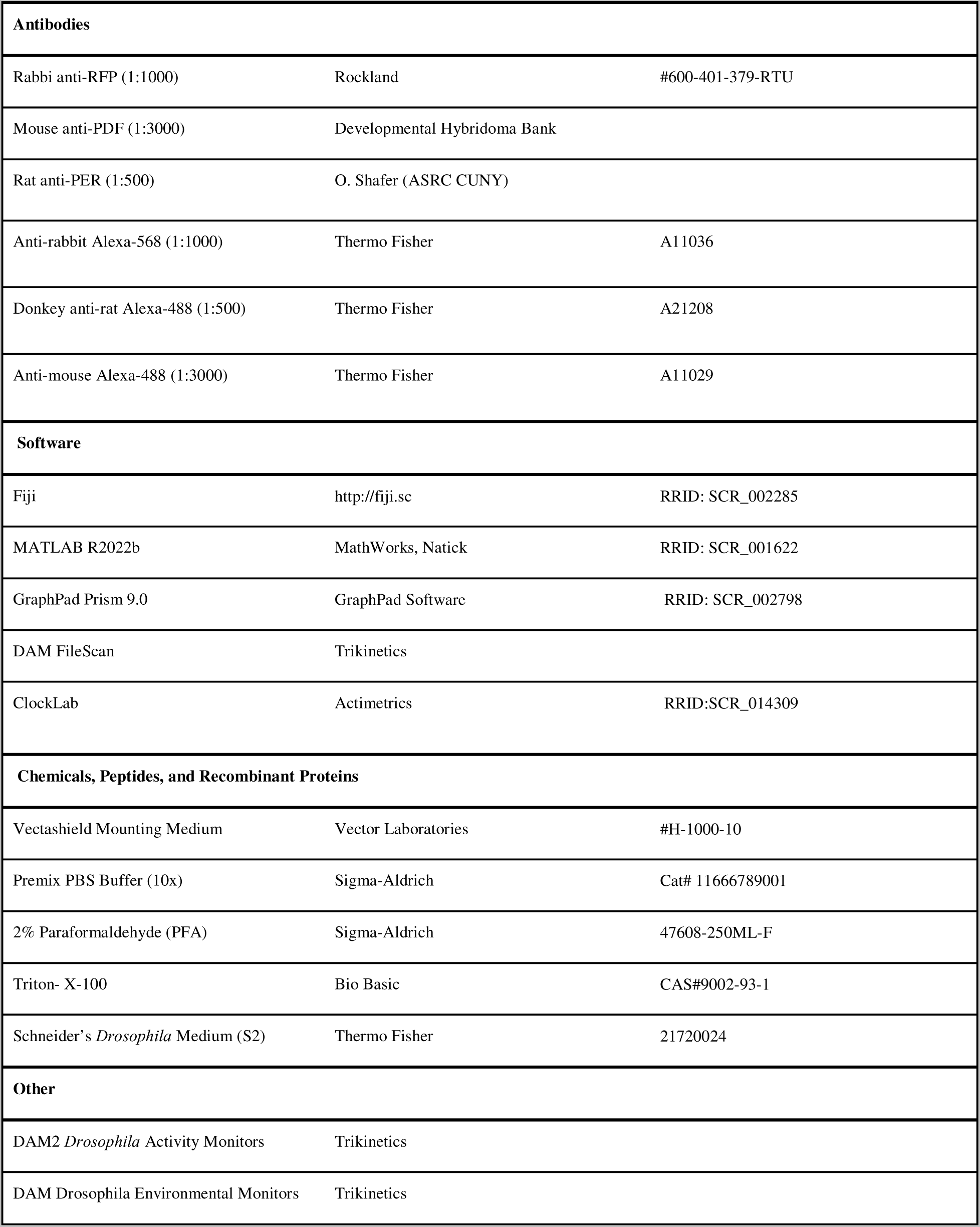

## Acknowledgments

We are very grateful to Justin Blau and Paul Hardin for their valuable feedback on various aspects of this project and for sharing fly lines with us. We are also grateful to M.Fernanda Ceriani, Amanda González-Segarra, Aishwarya Ramakrishnan Iyer, Orie Shafer, and Troy Shirangi for helpful comments on the manuscript, Annika Barber and Troy Shirangi for helpful discussions, and Orie Shafer for the rat anti-PER antibody. The mouse anti-PDF antibody was obtained from the Developmental Studies Hybridoma Bank, created by the NICHD of the NIH and maintained at The University of Iowa, Department of Biology, Iowa City, IA 52242. Stocks obtained from the Bloomington *Drosophila* Stock Center (NIH P40OD018537) were used in this study. This work was supported by an NSF Grant (IOS-2239994) to M.P.F.

---

**Supplementary Figure 1.**
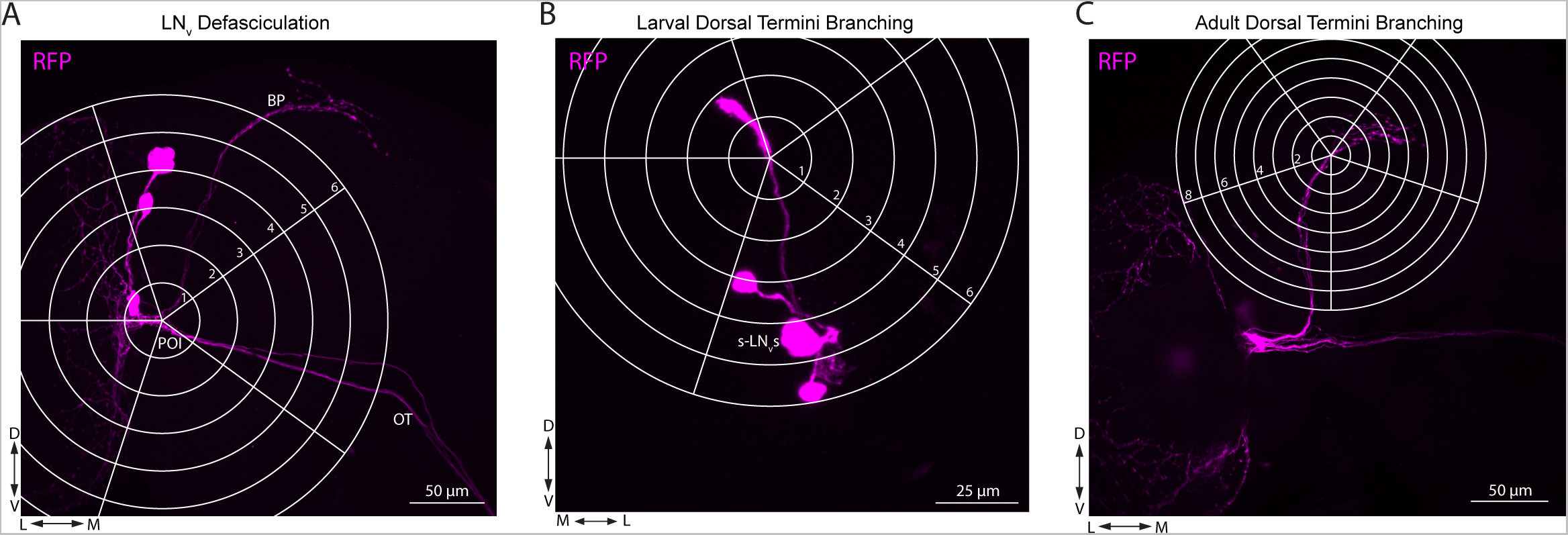
Quantification of defasciculation (ventral projections) and branching (dorsal projections) in the LNvs. (A-C) Representative confocal images of adult (A, C) and L3 larvae (B) control brains stained with anti-RFP (magenta). The sLN_v_s, OT, POI, and BP are indicated. Scale bars = 50 µm (A,C) and 25 µm (B). (A) To determine the degree of defasciculation of the sLN_v_ ventral projections in adult brains, 6 concentric circles separated by is 25 µm were centered at the point of intersection (POI), where the projections of the sLN_v_s and those of the lLN_v_s intersect. The most distant circle does not reach the main branching point (BP) in control brains; therefore, the dorsal termini are not included. The number of intersections between either the sLNvs or the lLN_v_s and each concentric circle were quantified. (B) Dorsal projection branching in the larval sLN_v_s was measured by counting the number of intersections the sLN_v_s had at each of the 6 concentric circles separated by 12.5 µm. (C) Adult sLN_v_ dorsal projection branching was measured by counting the number of intersections the sLN_v_s had at each of 8 concentric circles separated by 12.5 µm. This was adapted from a previous study ^27^ to capture the hyperextended projection phenotype of *Pdf* > Δ*-Clk* flies.

**Supplementary Figure 2.**
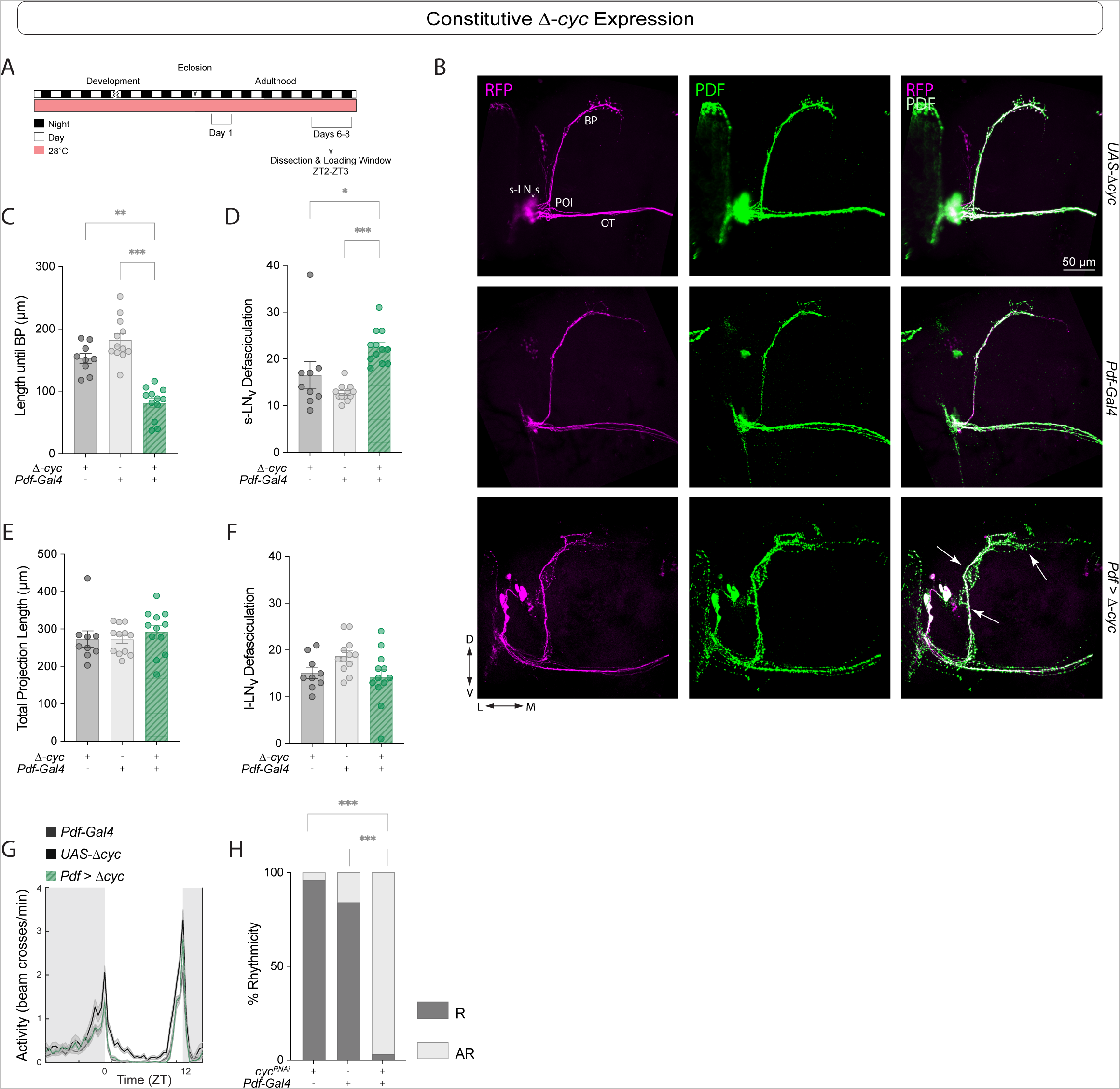
Δ-*cyc* expression in *Pdf*+ cells prevents sLN_v_s fasciculation. (A) Representative timeline of the experiments in the figure. Flies were kept in LD conditions at 28 °C for their entire lifespan. Dissections were performed within Days 6-8 post-eclosion from pupae at ZT2-3. (B) Representative brain confocal images of anti-PDF (green) and anti-RFP (magenta) staining in the sLN_v_s of flies in which Δ*-cyc* was constitutively expressed in the *Pdf*+ cells using a *Pdf-Gal4;tub-Gal80^ts^*driver. Each line also included a *Pdf-RFP* transgene. The images are representative of 2 independent experiments. Scale bar = 50 µm. Kruskal-Wallis tests followed by Dunn’s multiple comparisons tests were used to compare the sLN_v_ projection length until the branching point (C), the total number of intersections of the sLN_v_ ventral projections (D), the full sLN_v_ projection length (E), and the total number of lLN_v_ intersections (F). Each dot corresponds to one brain. For each genotype: 9 ≤ n ≤ 12. (G-H) Behavioral phenotypes of constitutive Δ-*cyc* expression. Experiments were conducted at 28°C. (G) Population Activity (left) plots for flies during days 3-5 of the LD cycle at 28°C (see Table 1 for additional quantifications). (H) Percent rhythmicity for the indicated genotypes under DD. R=Rhythmic and AR= arrhythmic. Fisher’s exact contingency tests were used to analyze the percentage of rhythmic flies under DD (DD1-8). * p < 0.05, ** p < 0.01, *** p < 0.001. Error bars indicate SEM. For each genotype: 24 ≤ n ≤ 32.

**Supplementary Figure 3.**
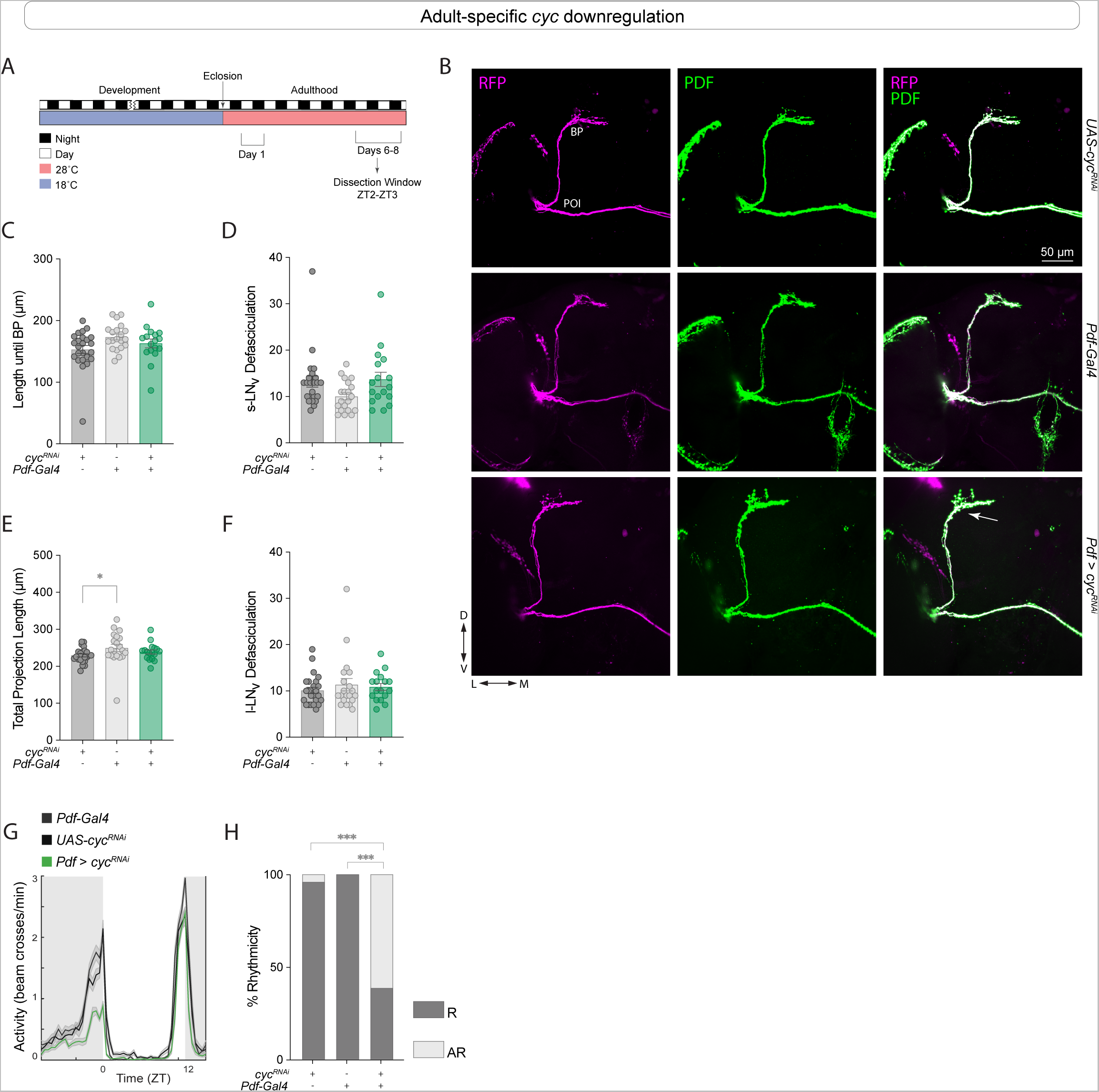
Adult-specific *cyc* knockdown does not affect sLNv neuronal morphology. (A) Representative timeline of the experiments in the figure. Flies were raised in LD at 18 °C, and transferred to 28 °C immediately after eclosion. Dissections were then performed within Days 6-8 post-eclosion from pupae at ZT2-3. (B) Representative confocal images of anti-PDF (green) and anti-RFP (magenta) staining in the sLN_v_s when *cyc* was downregulated exclusively after eclosion using a *Pdf-Gal4;tub-Gal80^ts^* driver. The images are representative of two independent experiments. Scale bar = 50 µm. Each line also included a *Pdf-RFP* transgene. Kruskal-Wallis tests followed by Dunn’s multiple comparisons tests were used to quantify the projection length until BP (C), the total number of intersections of the sLN_v_ ventral projections (D), the full sLN_v_ projection length (E), and the total number of lLN_v_ intersections (F). Each dot corresponds to one brain. For each genotype: 17 ≤ n ≤ 24. (G-H) Behavioral phenotypes of adult-specific *cyc* knockdown. Flies were raised in LD at 18 °C, before being transferred to 28°C upon eclosion. Experiments were conducted at 28°C. (G) Population Activity plots for flies during days 3-5 of the LD cycle at 18°C (see Table 1 for additional quantifications). (H) Percent rhythmicity for the indicated genotypes under DD. Fisher’s exact contingency tests were used to analyze the percentage of rhythmic flies under DD (DD1-8). * p < 0.05, *** p < 0.001. Error bars indicate SEM. For each genotype: 25 ≤ n ≤ 31.

**Supplementary Figure 4.**
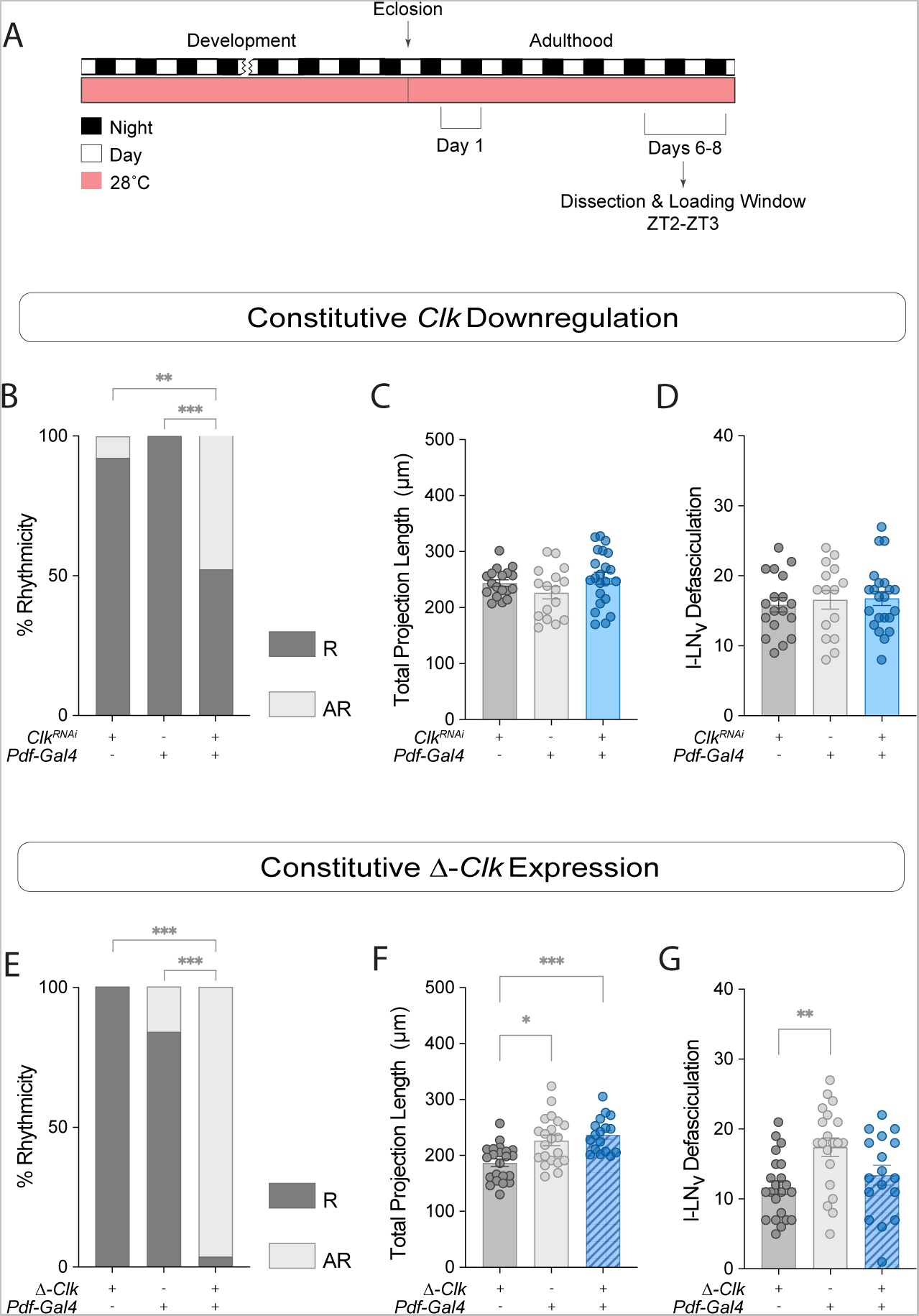
Expressing Δ*-Clk* in the LN_v_s results in morphology and behavioral phenotypes. (A) Representative timeline of the experiments in the figure. Flies were raised in LD at 28 °C For their entire lifespan. Behavioral assays and dissections were then performed within Days 6-8 post-eclosion from pupae at ZT2-3. Experiments were conducted at 28°C. (B) Fisher’s tests were used to compare the percent of rhythmic flies of each indicated genotype (additional quantifications can be found in Table 1). For each genotype: 21 ≤ n ≤ 26. (C-D) Additional quantifications of effects of *Clk^RNAi^*expression in the *Pdf*+ cells in adult brains. Neither the sLN_v_ total projection length (C) nor the lLN_v_ projections (D) were affected. Kruskal-Wallis tests followed by Dunn’s multiple comparisons tests were employed. For each genotype: 16 ≤ n ≤ 22. (E) Fisher’s tests were used to compare the percent of rhythmic flies of each indicated genotype (additional quantifications shown in Table 1). For each genotype: 27 ≤ n ≤ 31. (F-G) Effects of Δ*-Clk* expression in the *Pdf*+ cells in adult brains. The sLN_v_ total projection length (F) and the lLN_v_ projections (G) were quantified using Kruskal-Wallis tests followed by Dunn’s multiple comparisons tests. Each dot corresponds to one brain. For each genotype: 17 ≤ n ≤ 22. * P < 0.05, ** P < 0.01, *** P < 0.001. Error bars indicate SEM.

**Supplementary Figure 5.**
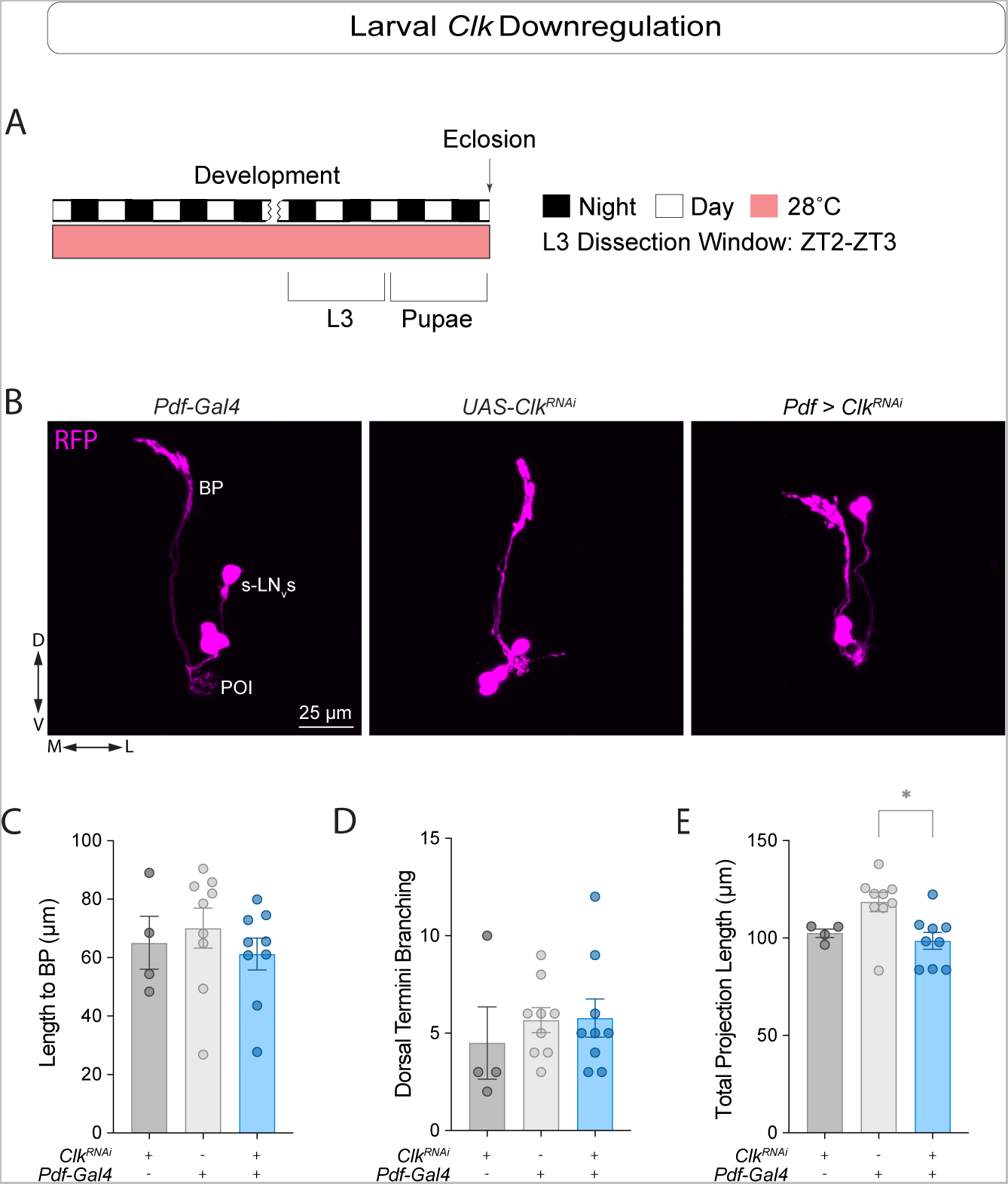
*Clk* downregulation in the larval sLN_v_s did not result in morphology phenotypes. (A) Representative timeline of the experiments in the figure. Larvae were raised in LD at 28 °C. Third instar larvae (L3) were dissected at ZT2-3. (B) Representative confocal images of anti-RFP (magenta) staining in the sLN_v_s when *Clk^RNAi^* was expressed in L3 larvae. The sLN_v_s, POI, and BP are labeled. Each line also contains a *Pdf-RFP* transgene. Scale bar = 25 µm. Kruskal-Wallis tests followed by Dunn’s multiple comparisons tests were used to compare the projection length from the POI to the BP (C), the total number of axonal intersections (D), and the total projection length from the POI (E). * p < 0.05. Error bars indicate SEM. Each dot corresponds to one brain. For each genotype: 4 ≤ n ≤ 9.

